# Screening and Identification of Azo Dye Decolourizers from Mangrove Rhizospheric soil

**DOI:** 10.1101/2022.02.24.481723

**Authors:** Akhilesh Modi, Sunita Singh, Jyoti Patki, Naveen Padmadas

**Affiliations:** Gujarat Biotechnology Research Centre, Sector 11, Gandhinagar, Gujarat India; School of Biotechnology and Bioinformatics, D.Y Patil deemed to be University, Navi Mumbai-400706 (India)

**Author notes:** **Corresponding Author:** Dr. Sunita Singh, Associate Professor, School of Biotechnology and Bioinformatics, D.Y Patil deemed to be University, Navi Mumbai- 400706 (India), **Email Address:**.

**Keywords:** Azo dyes, Bioremediation, Rhizospheric soil, Decolourization, Molecular studies, FTIR

## Abstract

Removal of synthetic textile dye poses a challenge to the textile industry and a threat to environment flora and fauna. These dyes are marginally degradable and recalcitrant, hence alternatives to physical and chemical techniques like various bioremediation studies involving plants, plant roots, single or consortium of microbes have been used as environment friendly methods for the removal of textile dye. In the present study potent bacteria for dye decolourization were isolated from rhizospheric soil collected from Kamothe, Navi Mumbai, India. Of the 20 isolates obtained after enrichment, seven isolates were used for further screening of efficient decolourization ability in MBM media containing 10% glucose, 2.5 % trace metal solution and 0.1% of MO dye concentration. Physiological parameters to optimize the decolourization of dye at optimum pH, temperature and incubation time was studied for all the seven isolates. UV-vis and Fourier Transform Infrared spectroscopy were used to investigate dye decolorization. The seven isolates were characterized morphologically, biochemically, and molecular identification of these strains were performed by 16S rRNA sequence analysis. The isolates were identified as *Bacillus paramycoides, Pseudomonas taiwanensis, Citrobacter murliniae, Acinetobacter pitti, Exiguobacterium acetylicum, Psychrobacter celer*, and *Aeromonas taiwanensis*. Out of these *Aeromonas taiwanensis* has shown exceptional capacity by 100% decolorization of azo dye in minimum time.

**Graphical Abstract:** 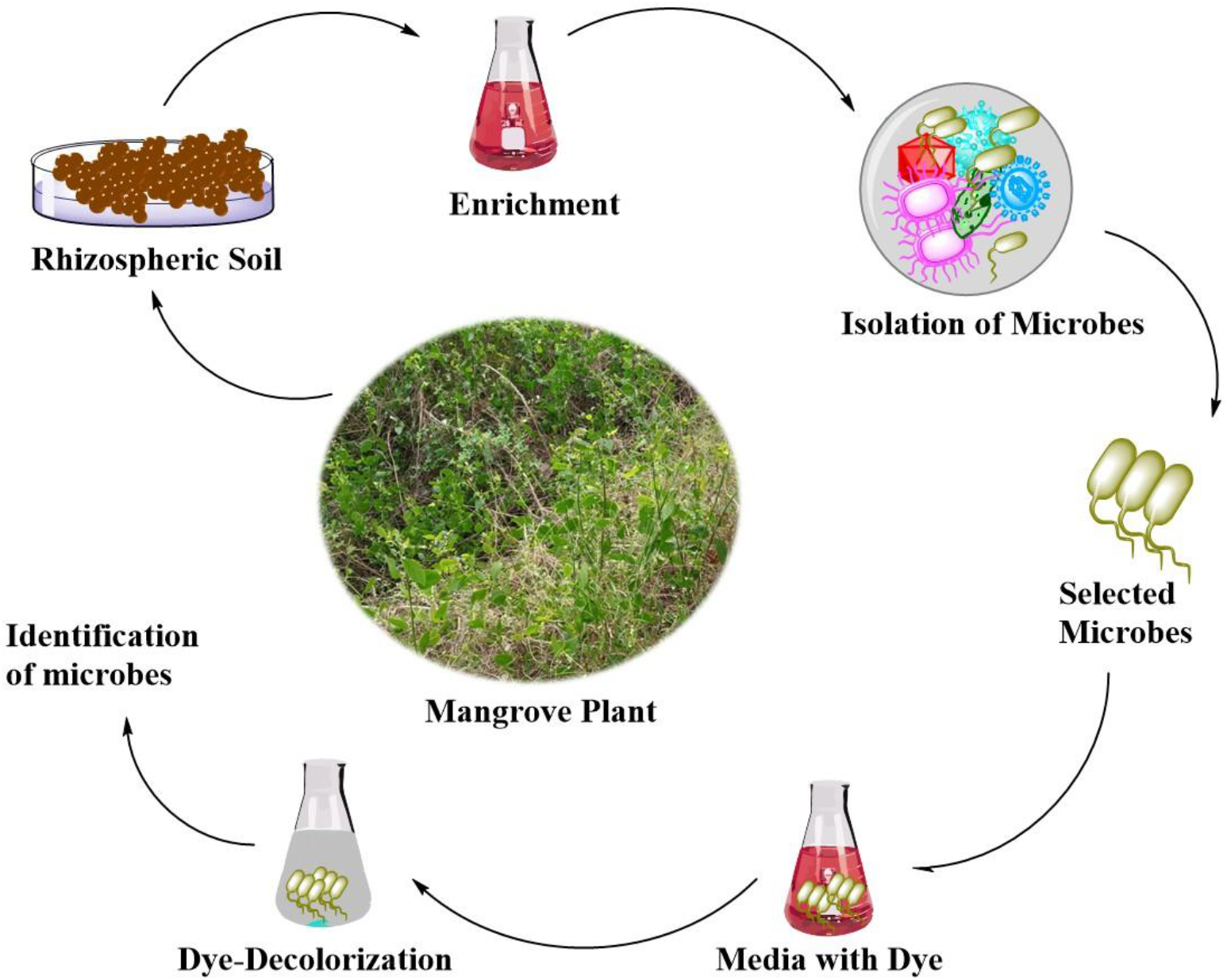

*Highlights of the study:*

1. Collection of soil samples from rhizospheric and non-rhizospheric region of the mangroves plant.
2. Enrichment of soil sample in media containing azo dye for dye decolourization.
3. Screening and isolation of potential dye decolourizers.
4. Optimize the condition for dye decolourization.
5. FTIR analysis and application based studies.
6. Identification of microbes using 16SrRNA.

## Introduction

Industrialization and overpopulation are the two main causes of pollution in the environment. The industrial, textile, transportation, and residential sectors are all expanding in sync with the population (Saravanan et al. 2018). Textile industries use dyes that are easy to make, less expensive, diverse, and colour-resolved. Synthetic dyes are extremely damaging to the environment when these businesses discharge their untreated waste product or effluent into fresh natural sources such as rivers, oceans, and ponds (Carliell et al. 1995; Zollinger et al. 2003). When industries release untreated toxins into the environment, they pollute the environment (Chiong et al. 2016). In many cases, the dye is broken down into by-products that are extremely poisonous, carcinogenic, and mutagenic to aquatic life (Weisburger 2002; Singh et al. 2017). Azo dyes, which are commonly used in the textile industry, are known to be hazardous to plants and animals (PS et al. 2013; Sudha et al. 2014). Various methods and procedures for waste water treatment have been devised in several countries, and all government rules and environmental agencies are on the lookout for the discharge of these industrial wastes. Biocolors generated from bacteria, fungi, and plants are now being researched as a possible replacement for synthetic dyes. When adequate raw materials are used, bio colours have various advantages over synthetic dyes, including reduced prices, faster extraction, higher yields, and no seasonal fluctuations. They are utilised as anticancer, anti-inflammatory, anti-microbial, and anti-oxidant agents, as well as food and cosmetic colouring additives. Isolation of novel microorganisms that create colour pigments could provide a new source of colorants for foods, textiles, and the pharmaceutical industry, among other applications (Heer and Somesh 2017). Colour and recalcitrant by-products have been removed from industrial waste water using physical, chemical, and combination approaches, as well as plant or microbe-based remediation strategies (Shaw et al. 2002). Physiochemical and chemical treatments, on the other hand, may have disadvantages such as high costs, large energy requirements, and the production of dangerous by-products. (Singh and Arora 2011). Currently, a microbial consortium of bacteria, fungi, algae, and yeast not only converts dye in waste water into simple substances, but also provides additional biomass (Chen et al. 2003; Shah et al. 2013). The studies on dye degradation has been reported among both anaerobes and aerobes like *Bacteroides* spp., *Eubacterium* spp., *Clostridium* spp, *Proteus vulgaris, Streptococcus faecalis, Bacillus* spp., *Sphingomonas* spp. and several yeasts. (Pearce et al. 2003; Forgacs et al. 2004; Lalnunhlimi and Veenagayathri 2016). Bacillus was shown to be dominant and decolorize the metanil yellow G (MYG) at temperatures ranging from 40 to 60ºC in a wide range of pH conditions in a study on the establishment of a halo-thermophilic Bacterial consortium for dye decolorization (Guo et al. 2021). Therefore, biological treatment for dye decolourization and degradation plays an important role in the cleaning of environment pollution (Abbas et al. 2019).

The discovery of novel bacterial strains for MO decolorization and degradation from pristine rhizospheric soil of mangroves forest was tried in this work. Various parameters have been studied in the meantime to better understand the performance of the strain in MO decolorization. The decolorization of dye was studied using UV– vis and Fourier transform infrared spectroscopy (FTIR). Furthermore, the microorganisms analysed in this study from the rhizospheric region would deliver the best results in the case of industrial use.

## Materials & Methods

### Collection of Soil sample

Rhizospheric (at depth of 10, 20 and 30 cm) and non-rhizospheric soil samples were collected from the pristine site of Kamothe in Navi Mumbai, India, as a source of dye decolorizing microbes. This region is defined by N latitude 19°10’0.4944” and E longitude 73° 5’47.2488”. Five soil samples were collected from the pristine site of mangroves plant using a sterile metallic spoon and sterile collection bags from the rhizospheric and non-rhizospheric sites. The adhering soil from the roots was carefully removed to isolate rhizospheric bacteria. Following collection, soil samples were kept at −20°C.

### Dyes & Chemicals

In this study, MO, MR and CR was used as a model dye, which was purchased from Himedia Labs Mumbai, India. The majority of the chemical and media components were obtained from Himedia Labs in Mumbai, India. All of the chemicals used in this study are of the highest purity and analytical grade. Unless otherwise specified, all media, buffers, solutions, reagents, micro centrifuge tubes, micro tips, glassware, and other items used in this study were sterilised at 15 lbs/inch^2^ for 20 minutes.

### Screening and Isolation of Dye decolourizing bacteria

Enrichment culture was used to isolate potent dye decolorizing bacterial strains. About 5 gm of rhizospheric and non-rhizospheric soil from each of the sampling area was added to the Nutrient broth (Peptone 5g/L, Sodium Chloride 5g/L, Beef extract 1.5 g/L, Yeast extract 1.5g/L) and Minimal Basal Medium (Disodium hydrogen phosphate 7g/L, Potassium hydrogen phosphate 3g/L, Sodium citrate 0.5g/L, Magnesium sulphate 0.1g/L, Ammonium sulphate 1g/L) containing 10% glucose solution and trace element (Ferrous sulphate 0.5g/L, Zinc sulphate 0.5g/L, Manganese sulphate 0.5g/L, 0.1N H_2_SO_4_ 10ml/l) supplemented with 0.1 % (100 mg) and 0.2 % (200 mg) of MO dye concentrations and incubated at room temperature under static conditions. The amount of dye decolorization was measured every 6 hours until the incubation time was 48 hours. A small amount of the decolorized medium was streaked on Nutrient Agar containing MO dye. After 48 hours of incubation at 37°C, organisms that showed a zone of decolorization were chosen and subsequently screened based on their dye decolorization potential. A1, A2, M41, M42, M43, M3, and M53 were the strains that showed the most decolorization and were chosen for future study.

### Experiment of dye decolourization

The decolorization of MO dye was studied using the batch culture method. The entire experiment was carried out in triplicate. The isolates from rhizospheric and non-rhizospheric soil that showed maximum decolorization zone on Nutrient agar plate and MBM agar plate supplemented with 0.1 % MO dye were grown on both MBM and NB media (pH 7.0) and incubated in static condition at 37°C for 24 hours to obtain synchrony in the growth phase (i.e. late exponential or early stationary phase). All the seven isolates were used at a constant density (0.1OD at 600nm) for further research.

### Effect of dye concentration on decolourization

In order to study the initial influence of dye on decolorization in static conditions at various time intervals, only two distinct concentrations of dye were studied in the decolorizing medium viz. 0.1% (1000mg/L) and 0.2% (2000mg/L).

### Effect of bacterial isolate and Cell Free Extract on dye decolourization

The effect of both the single bacterial isolate and its cell free extract (CFE) on dye decolorization was investigated. The overnight grown bacterial culture was added to freshly made decolourizing media containing 0.1% (1000 mg/L) MO dye concentration and cultured at 37°C under static condition for 48-72 hours to evaluate the dye decolorization. In the case of CFE analysis, the bacterial culture was cultured in decolorizing media for 24 hours and centrifuged for 10 minutes at 10,000 rpm. To monitor dye decolorization, the supernatant (CFE) was added in a 1:10 ratio to freshly made decolorizing medium containing 0.1 % (1000mg/L) dye concentration and incubated at 37°C in a static environment for 48-72 hours.

### Effect of different media on dye decolourization

The ability of isolated microorganisms to decolorize MO dye was tested only under static conditions. For dye decolorization, a general purpose medium such as NB and a minimal media such as MBM were used. MO dye decolorization was performed in nutrient broth medium and MBM medium with 10% glucose and trace element solution containing 0.1 % dye concentration. Each bacterial isolate’s CFE was added to NB medium and MBM medium containing 0.1 % dye and incubated at 37°C in static condition for 48-72 hours to monitor dye decolorization.

### Effect of incubation time on dye decolourization

CFE of the respective isolate was added individually in a ratio of (1:10) in a fresh MBM dye decolourizing medium and incubated at 37°C. To verify for abiotic dye decolorization, an untreated control sample was incubated under the same conditions. Samples were taken at 2-hour intervals and spectrophotometric analysis (at 463 nm) was used to check for dye decolorization.

### Effect of pH on dye decolourization

The influence of pH on dye decolorization was investigated in MBM medium supplemented with 0.1 % MO dye at pH values ranging from 6 to 8.5. (6, 7 and 8.5). In a freshly prepared dye decolorizing medium, the CFE of a respective isolate was added and incubated at 37°C. The untreated control sample was also incubated under the same conditions to test for abiotic dye decolorization. Samples were taken at 2-hour intervals and spectrophotometrically studied for dye decolorization.

### Effect of temperature on dye decolourization

The effect of temperature on MO dye decolorization was investigated at R.T and 37°C. The CFE was prepared and added into the dye decolorizing medium. Dye decolorization was observed at 2 hour intervals for up to 66 hours.

### Decolourization of mixture of Azo dyes by bacterial monoculture

The mixture of three distinct azo dyes consisted of MO, MR, and CR (λ max in the range of 400-500 nm). An azo dye mixture’s absorbance peak (λ max) value was found to be at 455 nm. The decolorization of dye mixtures was investigated by inoculating CFE of M53 isolate (most prominent isolate) in MBM medium supplemented with 10% glucose and trace element solution and incubating it under static conditions at 37°C. The un-inoculated control sample was also incubated under the same conditions to test for abiotic dye decolorization. The samples were taken out at 2 hour intervals and tested for dye decolorization at 455 nm.

### Analytical experiment

#### Spectrophotometric analysis of dye decolourization

UV-VIS spectrophotometer analysis was used to evaluate dye decolorization. The absorbance peak (λ max) value of MO in decolorizing media was obtained by scanning the absorption of light within the visible range (400-700nm) at 10 nm intervals using a double beam UV-Vis spectrophotometer. The wavelength with the highest optical density was considered to be the maximum wavelength (λ max). As a result of this process, the maximum of MO was determined and found to be at 463 nm. The %age of decolorization was estimated by subtracting the starting and final absorbance readings.

A sample of culture broth (1 ml) was centrifuged for 10 minutes at 10,000 rpm. To avoid media influence, the absorbance of supernatant was measured against a blank and plotted against incubation time in Microsoft® office excel 2016 using the “scatter” function, a trend line was added, and the R^2^ value was computed. The following formula was used to compute the %age of decolorization

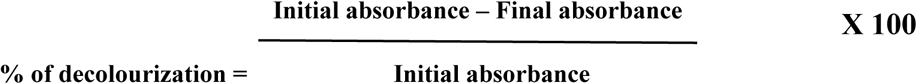

### Analysis of bacterial growth

The growth of all isolated bacterial cultures was spectrophotometrically monitored at 600nm at varied time intervals during dye decolorization **(Fig S1)**.

### FTIR analysis of decolourized product of azo dye

FTIR analysis was performed to check the dye decolourization. After complete decolorization, the decolorized liquid was centrifuged at 10,000 rpm for 15 minutes to get the supernatant. The FTIR analysis of CFE with dye before and after treatment was performed and the spectra was compared to a control dye in the mid IR range of 500-4000 cm-^1^ with a scan speed of 16 scans. In a 1:200 ratios, the sample was combined with spectroscopically pure KBr.

### Statistical analysis

All experiments were carried out in triplicate, and mean SD data were reported. Microsoft Excel was used to do the statistical analysis (version 2016). To ensure reproducibility of results, data was analysed using the standard deviation and portrayed as an error bar, as in all figure presentations. Microsoft Excel (version 2016) was used to perform ANOVA (single factor) for all isolates at their respective pH and temperature following complete dye decolorization.

### Identification of dye decolourizing bacteria based on 16S rRNA partial sequence analysis

Sanger sequencing was used to identify bacterial strains having high dye tolerance. For 16S rRNA identification, the bacterial strains were sent to the National Centre for Microbial Resources-NCCS in Pune, India.

### Phylogenetic analysis

The nucleotide sequence was first analysed at the NCBI server’s Blast-n site (http://www.ncbi.nlm.nih.gov/BLAST) and the associated sequence was downloaded. MEGA 11 software was used to align the sequence (Tamura et al. 2021). The Neighbour-Joining approach was used to infer evolutionary history (Saitou and Nei 1987). Next to the branches is the %age of duplicate trees in which the related taxa clustered together in the bootstrap test (1000 replicates) (Felsenstein 1985). To convert distance to time, a clock calibration of 0.01 (time node-1 height) was employed. The evolutionary distances were computed using the Maximum Composite Likelihood method (Tamura et al. 2021). The phylogenetic tree was constructed using the aligned sequences using MEGA 11 and designing of the tree was done using the Interactive Tree Of Life (https://itol.embl.de) is an online tool (Letunic et al. 2021).

### Application based study for dye decolorization of MO

A sterile white cotton fabric measuring 2cm by 2cm was dyed with 0.1 % MO and dried for 4 hours at 60°C. Following the drying of the fabric, 100 ml of MBM media enriched with 10% glucose and trace element solution and CFE of M53 isolate was applied and incubated at 37°C in a static environment. To check for abiotic decolorization of MO dye, the fabric was placed in sterile medium containing MO dye (control) and incubated under the same conditions. Dye decolorization was seen at both the visual and spectrophotometric levels (fabric and the medium of test and control).

## Results

### Screening & Isolation of dye decolourizing bacteria

Dye-decolorizing bacteria were isolated from rhizospheric and non-rhizospheric soil based on their ability to form a clear zone on nutrient agar plate containing MO (**Fig S2**). A large number of bacterial isolates were obtained and screened further to identify those with the best decolorization ability. Following multiple screening, seven bacterial isolates from rhizospheric soil viz. A1, A2, M41, M42, M43, M53 and M3 isolate from non-rhizospheric soil were used for further decolorization experiments.

### Effect of dye concentration on decolourization

Soil samples were acclimatised to media containing 0.2 % MO. Although cell growth was seen, there was less extent of decolorization **(Fig S3A)**. The efficiency of the soil microbes for decolorizing the model dye (MO) at concentrations ranging from (1000mg-2000mg/L) was analysed and for further studies 0.1 % of dye was used. All isolates obtained the highest decolorization with a dye dosage of 1000 mg/L **(Fig S3B)**.

### Effect of media on dye decolourization

On comparing the effect of media on dye decolorization, MBM medium was found to exhibit decolorization because isolates in NB media generated black pigments, resulting in difficulties for analysis of MO dye decolorization. Whereas, in the case of MBM, greater dye decolorization was achieved and analysis was simpler without any interference of media blackening (**Fig S4)**.

### Effect of bacterial culture & CFE on dye decolourization

Decolorization of dye in the presence of cell (O.D = 0.1) and CFE (after 24 hours) in decolourization medium was compared. After 48 hours, the former trial resulted in black pigmentation and medium darkening in contrast to the control without blackening **(Fig 1A)**. CFE collected after 24 hours was applied and monitored for 48-72 hours to provide better MO dye decolorization than the culture itself **(Fig 1B)**. As a result, all further research was conducted using respective CFE of grown isolate in MBM medium supplemented with 10% glucose and a trace element solution.

**Figure 1(A).**
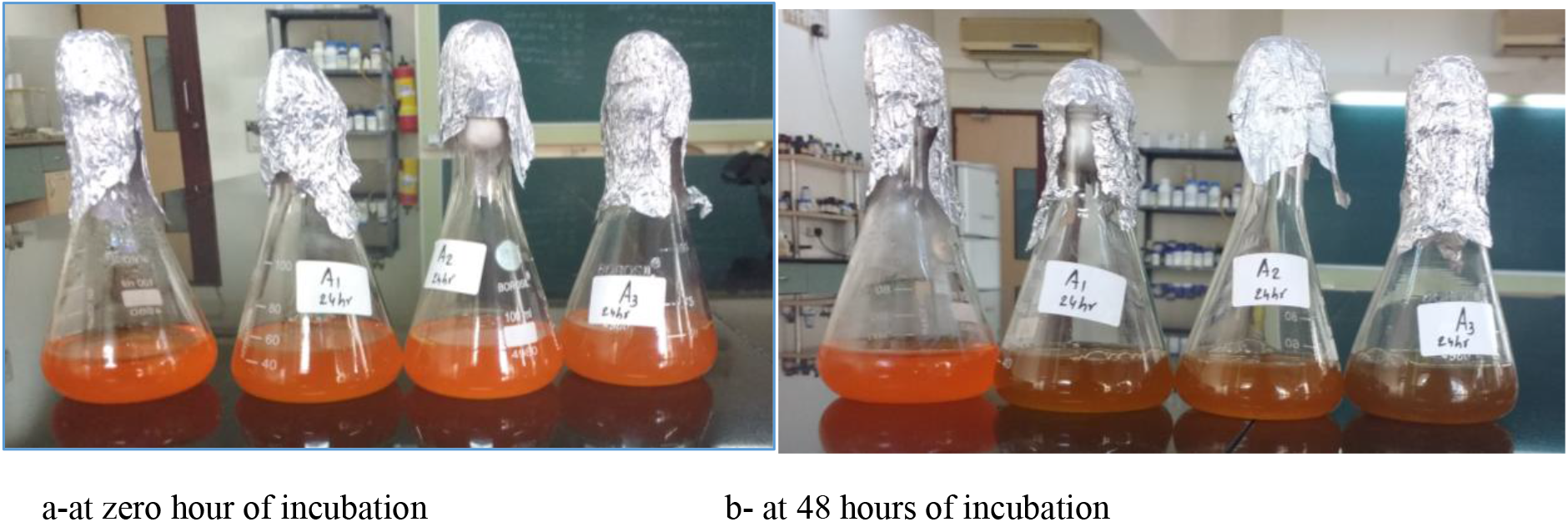
Effect of bacterial culture & CFE on dye decolourization. Bacterial culture isolates in MBM media with 0.1% Methyl Orange dye before and after incubation at 37°C

**Figure 1(B).**
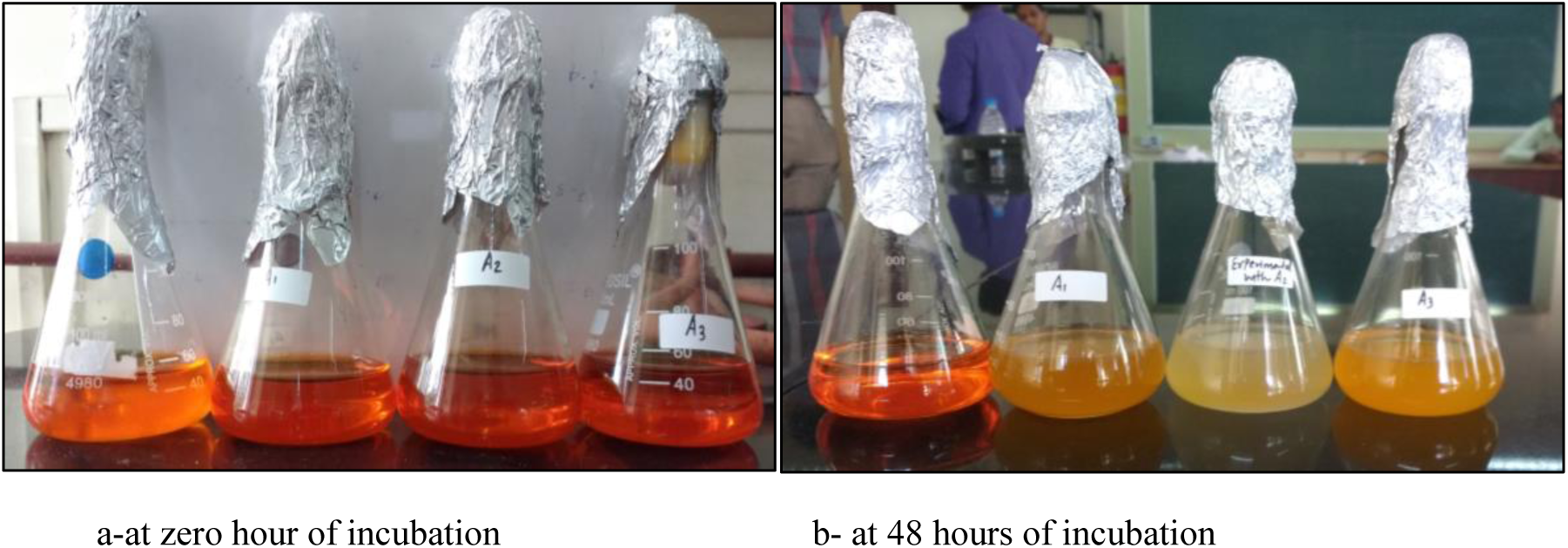
Cell Free Extract of isolates in MBM media with 0.1% Methyl Orange dye before and after incubation at 37°C.

### Effect of incubation time on dye decolourization

The results showed that bacterial isolates from rhizospheric soil viz. A1, A2, M41, M42, and M43 showed 82.2, 81.4, 93.3, 95, and 88.1 % decolorization of 0.1 % MO dye after 66 hours at 37°C in static condition. Isolate M53, on the other hand, achieved nearly 100% dye decolorization in just 26 hours. In contrast, an M3 isolate from non-rhizospheric soil was able to achieve 95% decolorization **(Fig 2)**.

**Figure 2.**
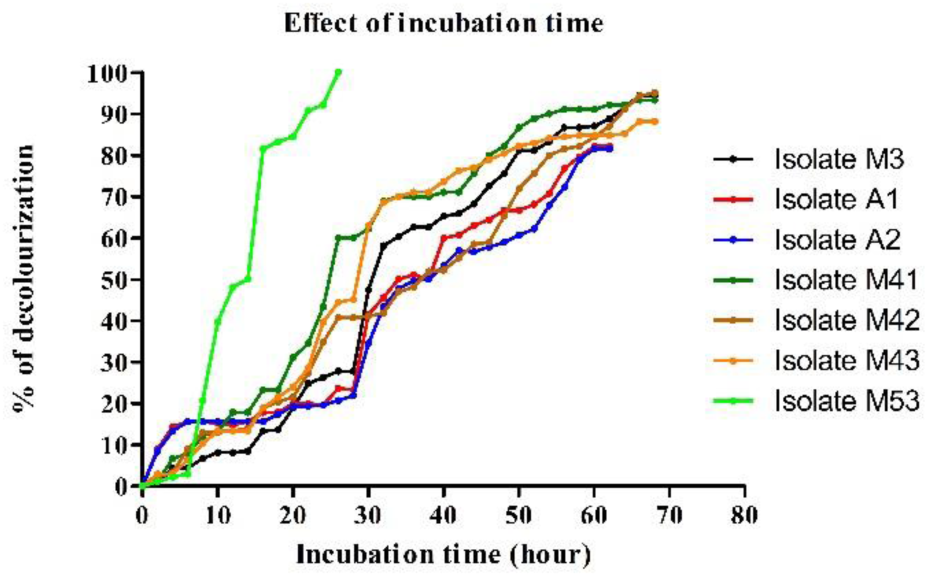
Effect of incubation time on dye decolourization. Effect of incubation time on decolourization of Methyl Orange by bacterial isolate A1, A2, M41, M42, M43, M53 & M3 (at 1000 mg/L dye concentration, pH 7.0, 37°C under static condition after 66 hours of incubation given as mean ± SD)

### Effect of pH on dye decolourization

The study looked at the growth and decolorization abilities of the seven isolates at pH 6, 7, and 8.5. Except for the M3 isolate, which did not grow at pH 6 or pH 8.5 at 37°C, all of the pH levels permitted the isolates to grow. For all isolates, maximum dye decolourization was reported at pH 7.0. **(Fig 3b)**. Bacterial isolates decolorize MO dye much less at pH 6.0 **(Fig 3a)**. For all isolates except M53 isolate yielded about 100% dye decolorization at all pH ranges. **(Fig 3 & Fig S5)**.

**Figure 3.**
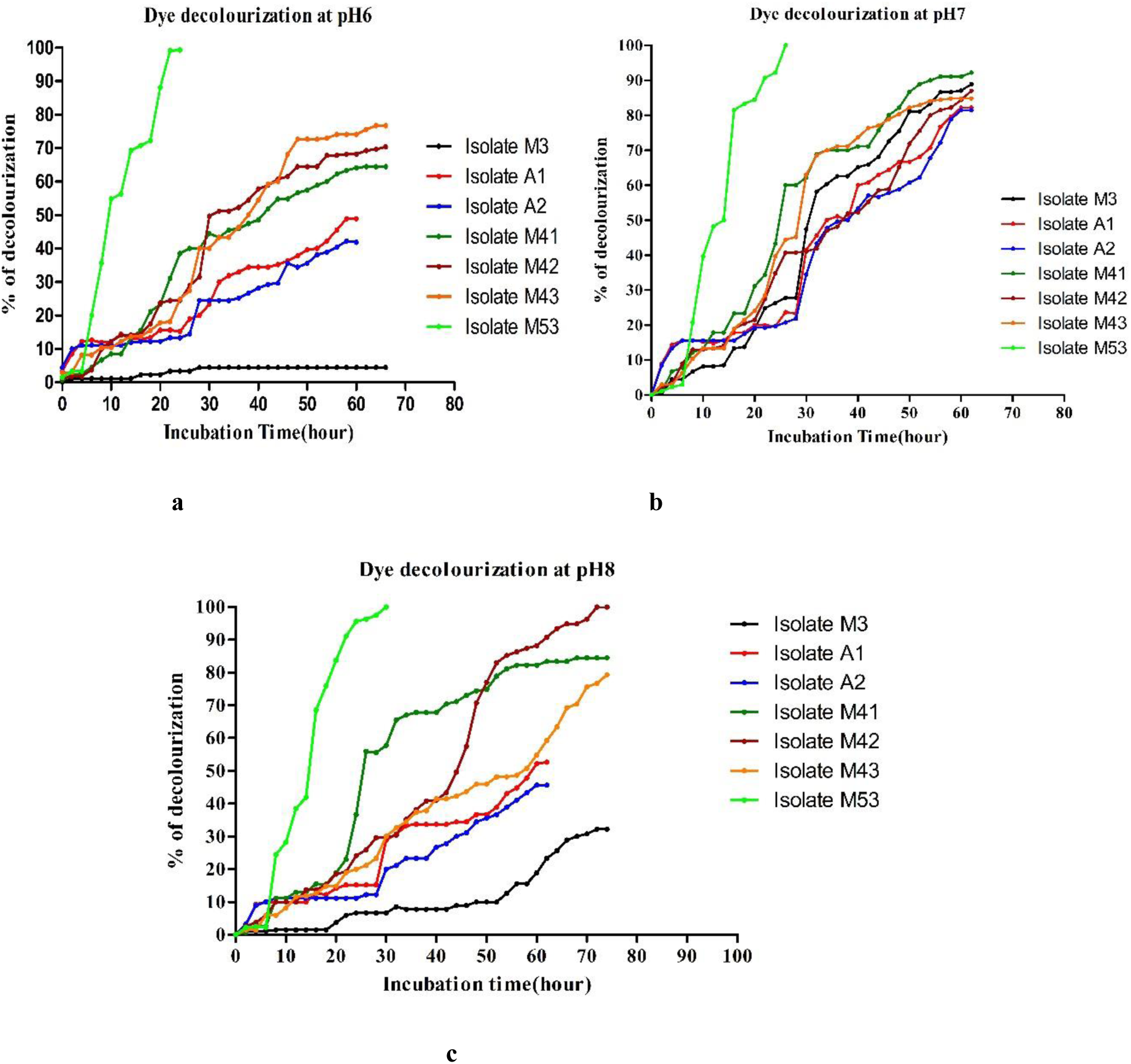
Effect of pH on dye decolourization. Effect of pH on decolourization of Methyl Orange by bacterial isolate A1, A2, M41, M42, M43, M53 & M3 (at 1000 mg/L dye concentration, pH 6, 7 and 8.5, 37°C under static condition after 66 hours of incubation given as mean ± SD)

### Effect of temperature on dye decolourization

In the present, study two temperature conditions like 25°C and 37°C for all 7 bacterial isolates were tested. The maximum decolourization of MO dye was observed at 37°C for all isolated cultures **(Fig S6)**. With further decrease in temperature to the 25°C to 27°C, decolourization of MO dye was drastically decreased. The %age of decolourization of MO increased at temp range from 25°C (8.9%) to 37°C (∼100%). These results indicate an optimum temperature for decolourization of MO dye was at 37°C for all the isolates. At 25°C temp M41, M42 and M53 showed more than 90% of decolourization of MO **(Fig 4)**., Indicating its application under various environmental conditions.

**Figure 4.**
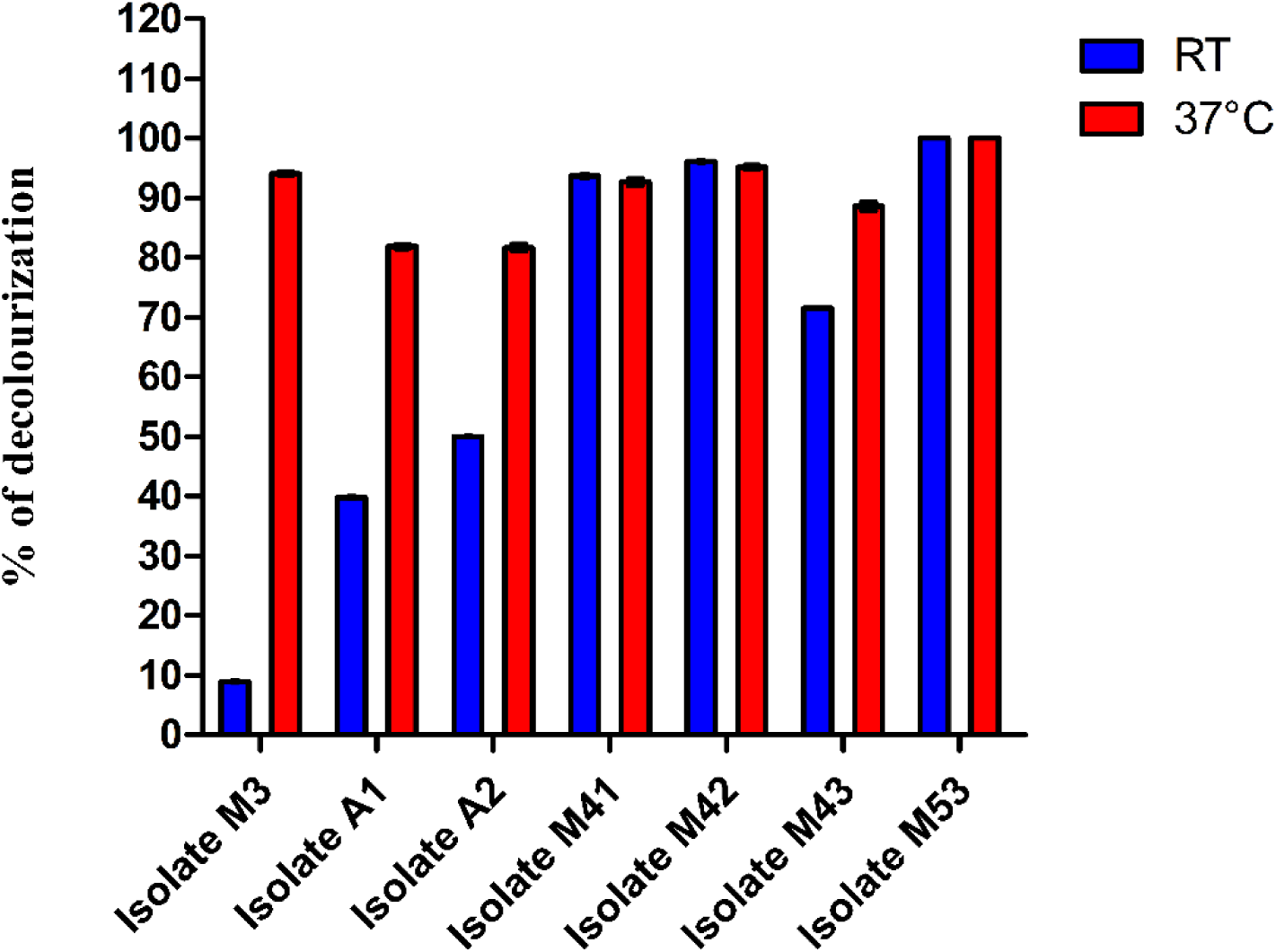
Effect of temperature on dye decolourization. Effect of Temperature on decolourization of Methyl Orange by bacterial isolate A1, A2, M41, M42, M43, M53 & M3 (at 1000 mg/L dye concentration, pH 7.0, under static condition after 66 hours of incubation given as mean ± SD)

### Decolourization of Azo dyes mixture by bacterial monoculture

Only one bacterial isolate M53 was tested in MBM media with 10% glucose and trace element solution supplemented with 0.1 % azo dye mixture (MO, Methyl Red, Congo Red) and incubated at 37°C under static condition for 48 hours **(Fig 5A)**, and optimum (∼100%) decolorization **(Fig 5B)** was observed after 48 hours of incubation at 37°C under static condition.

**Figure 5(A).**
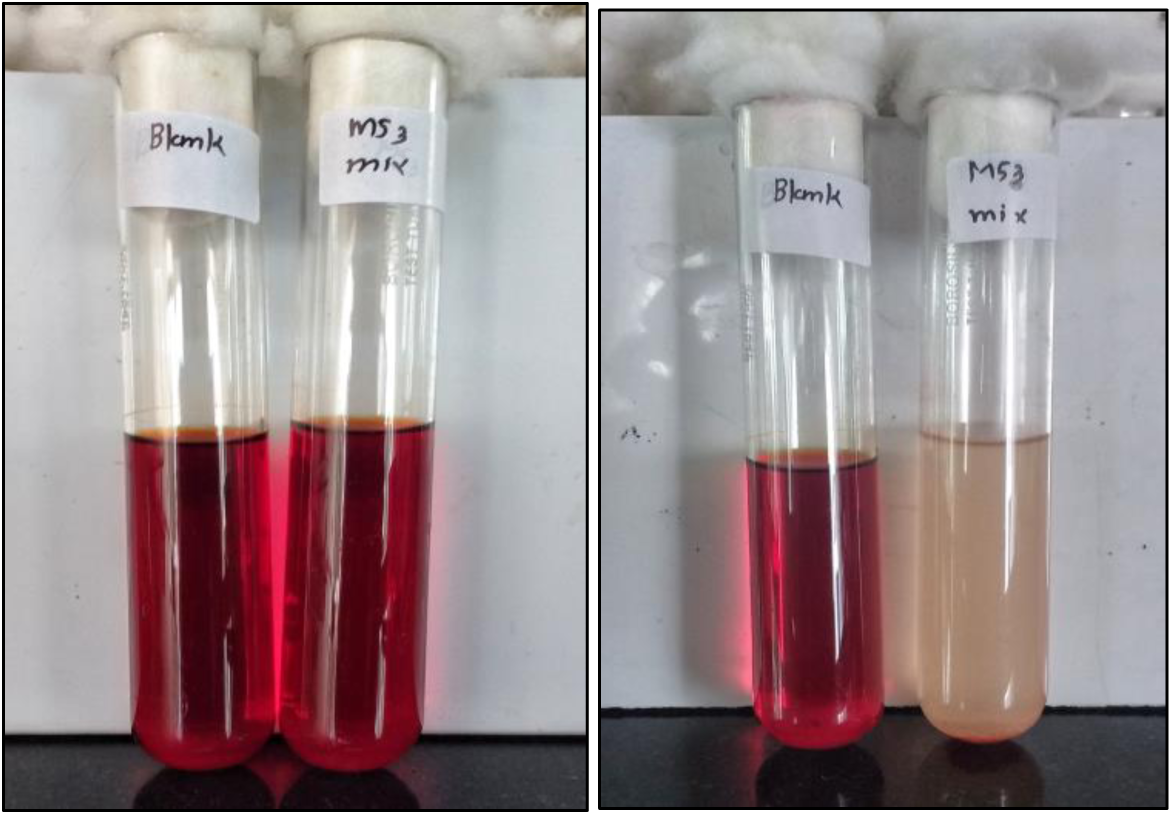
Decolourization of Azo dyes mixture by bacterial monoculture. Decolourization of Mixture of azo dyes by M53 isolate (at 1000 mg/L dye concentration, pH 7.0, 37°C under static condition)

**Figure 5(B).**
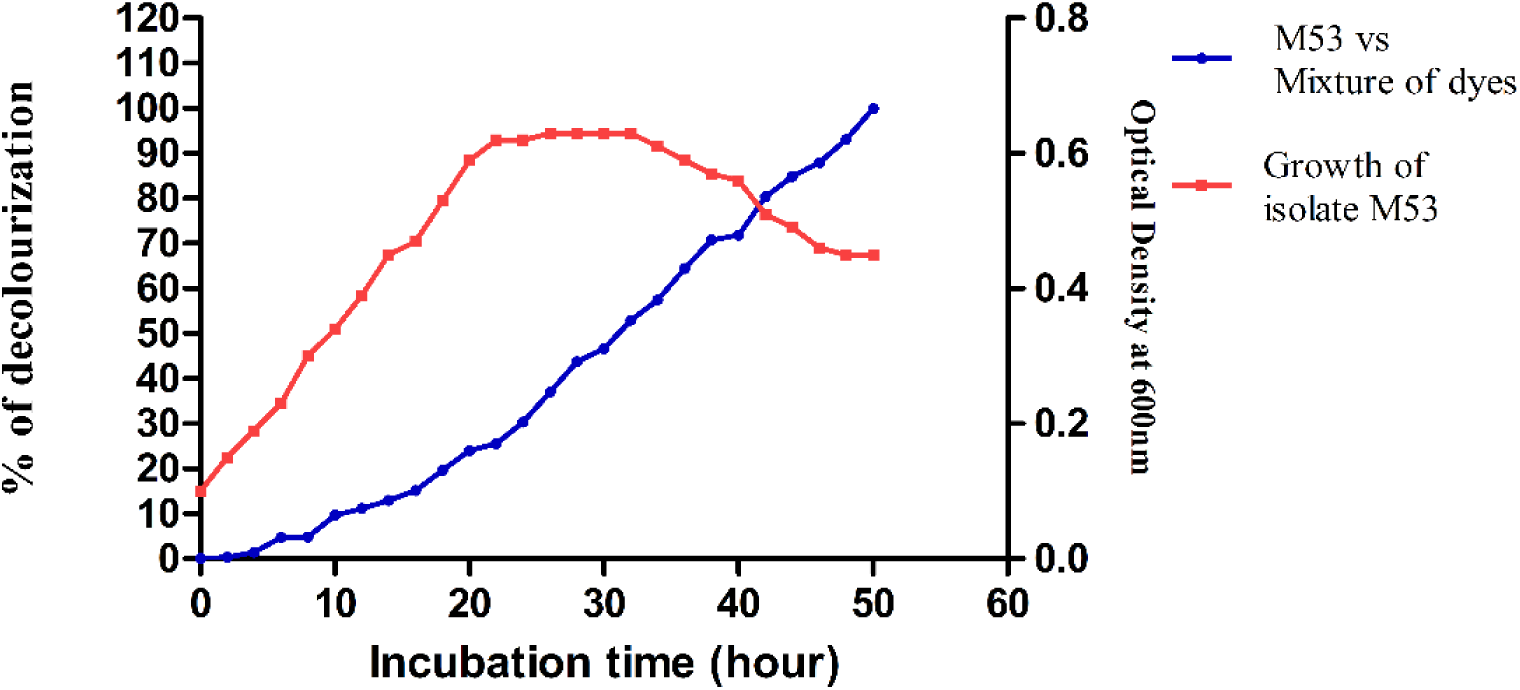
Decolourization of Mixture of azo dyes by M53 isolate (at 1000 mg/L dye concentration, pH 7.0, 37°C under static condition given as mean ±SD)

### FTIR analysis of decolourized product of MO dye

The FTIR spectra of untreated dye versus decolorized dye product is depicted in **(Fig 6)**. Peak at 1597.11 cm-1 in the case of untreated dye, which is indicative of –C=C-aromatic skeletal vibrations. Peak at 1114.89 cm-1 indicates –S=O vibration, while peak at 1313.57 cm-1 indicates C=N vibration. CO, CN, or phenolic vibration is assigned to 1026.16 cm-1,1114.89 cm-1,1159.26 cm-1,1003.02 cm-1. Asymmetric -CH2 vibration is attributed to peak value 1410.01cm-1. All of the peak values were missing in the biodecolorized product **(Fig 6)**, indicating that the linked groups on the benzene ring structure were stretched. After 48 hours, the FTIR spectrum of the decolourized product by M53 bacterial isolate shows new peaks at 1633.76 cm-1 and 1078.24 cm-1, and after 72 hours, new peaks at 1631.83 cm-1 and 1072.46 cm-1, which can be attributed to phenolic vibrations –C=O and C=O, C=N.

**Figure 6.**
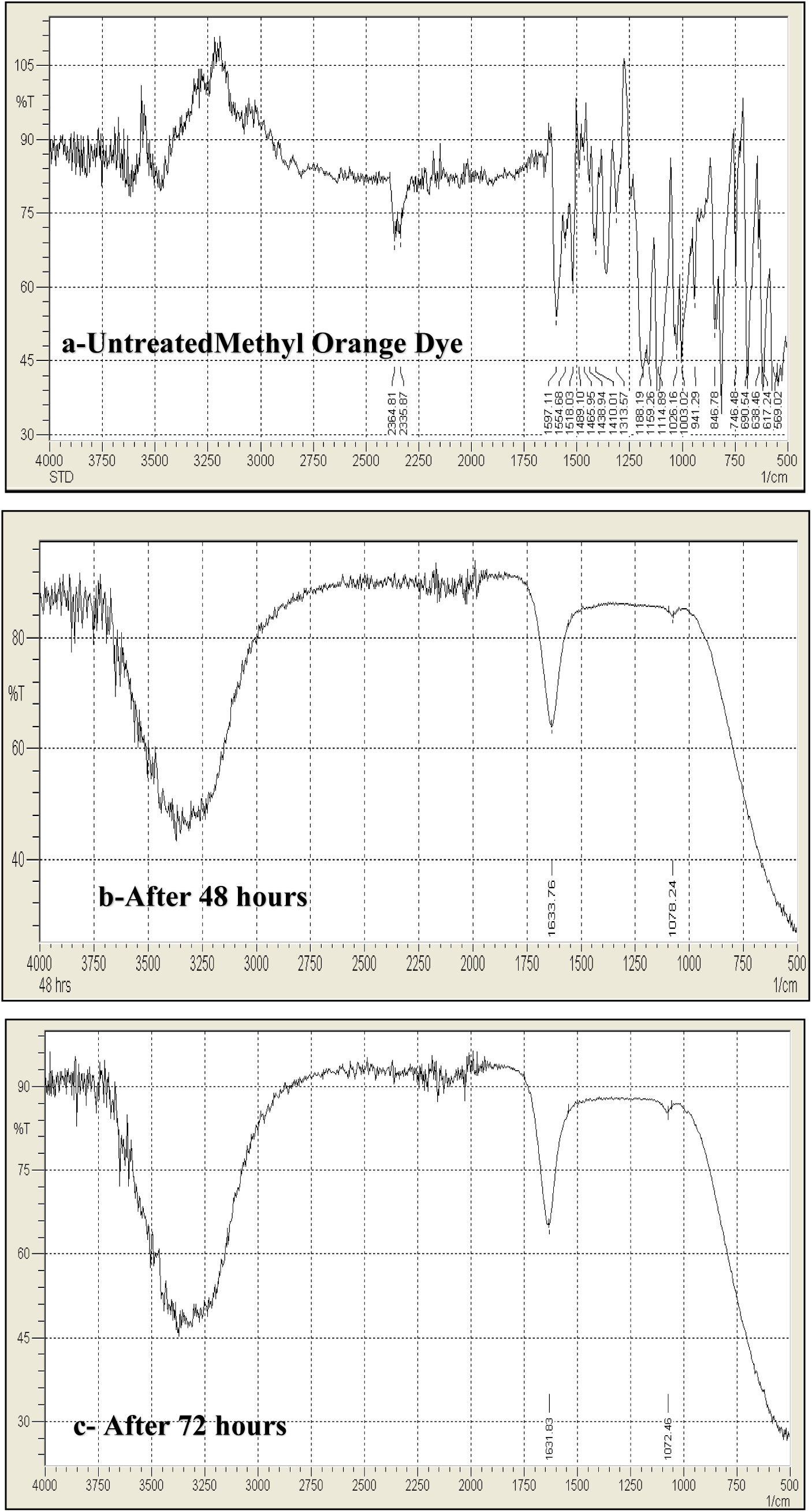
FTIR analysis of decolourized product of Methyl Orange dye. FTIR spectrum analysis of Methyl Orange dye and its biodegraded/biodecolorized products

### Statistical Analysis

**Table S2**. shows the findings of the one-way ANOVA. P values were 3.73 × 10^−44^, 8.27 × 10^−45^, and 1.18 × 10^−52^ at pH 6.0, 7.0, and 8.5, respectively, which were less than 0.05, while at 25°C and 37°C, p values were 6.3 × 10^−44^ and 8.27 × 10^−45^, which were also less than 0.05. All isolated samples had F values that were higher than the F-critical levels. As a result, at a 95 % confidence interval, the means of each set are considerably different from one another.

### Identification based on 16S rRNA sequence analysis

For all isolates, the partial 16S rRNA resulted in a nucleotide length of 700bp. MEGA 11 software was used to perform the phylogenetic analysis of the sequences. The trimmed, combined 16S sequences from each isolate were compared against all genomic databases using the web application Megablast with the default parameters (Morgulis et al. 2008). The Mega Blast result was also confirmed using EzBioCloud-EzTaxon, a web-based tool for identifying prokaryotes based on 16S ribosomal RNA gene sequences (Yoon et al. 2017). The isolates A1, A2, M41, M42, M43, M3 and M53 were identified as *Bacillus paramycoides, Pseudomonas taiwanensis, Citrobacter murliniae, Acinetobacter pitti, Exiguobacterium acetylicum, Psychrobacter celer*, and *Aeromonas taiwanensis* **(Fig 7)**. All isolates’ sequences have been submitted to the genbank under the accession numbers MT256266 (A1 isolate), MT256267 (A2 isolate), MT256269 (M41 isolate), MT256270 (M42 isolate), MT256271 (M43 isolate), MT256273 (M3 isolate), and MT256274 (M53 isolate) **(Table S1)**.

**Figure 7.**
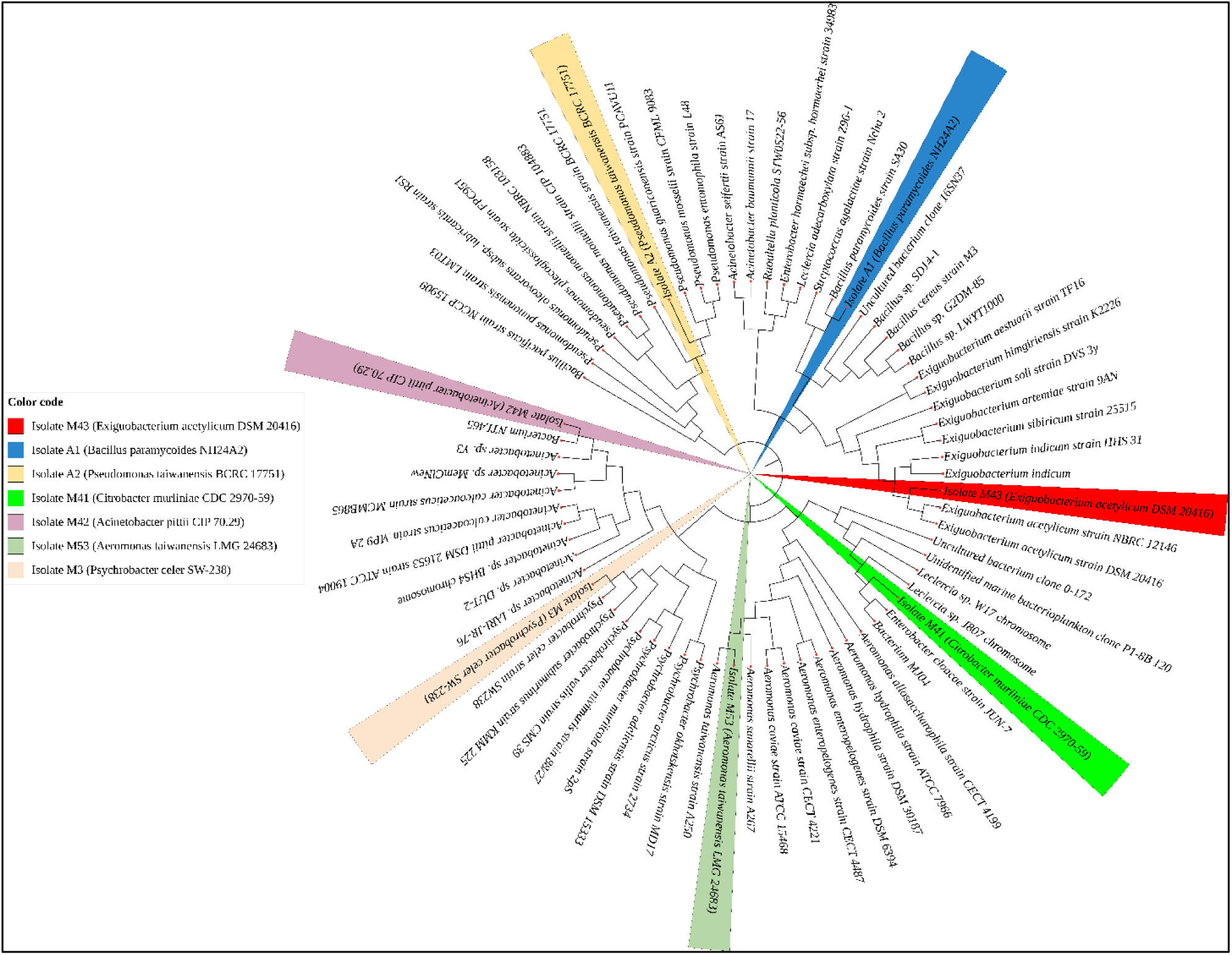
Identification based on 16S rRNA sequence analysis. Phylogenetic tree analysis of seven isolates for dye decolourization

### Application based study for dye decolourization of MO

In order to evaluate the efficiency of the dye removal on the cloth, the CFE of the most efficient isolate was subjected to further analysis. Coloured fabric was incubated at 37°C under static conditions in MBM media containing 10% glucose and trace element solution, which was supplemented with CFE of M53 isolate. Visual decolorization of fabric was recorded every 2 hours, while spectrophotometric decolorization of decolorizing fluid was detected every 2 hours. After 21 hours of incubation at 37°C in a static setting, the coloured cloth had completely decolourized **(Fig 8)**.

**Figure 8.**
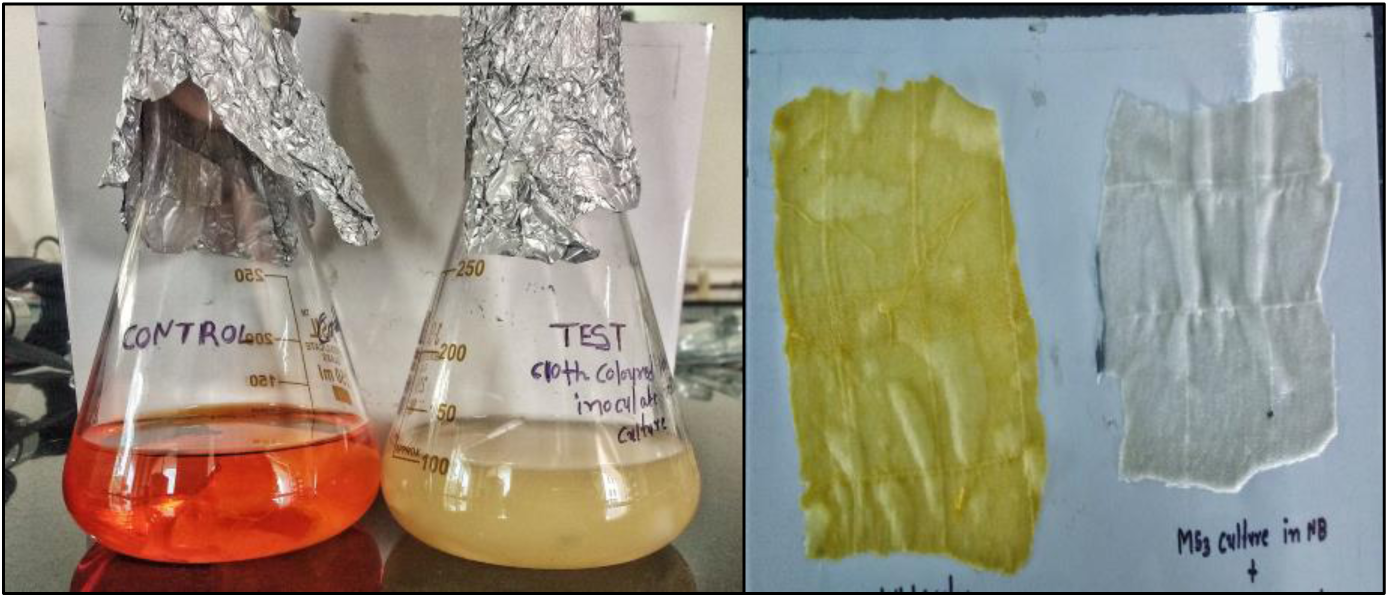
Application based study for dye decolourization of Methyl Orange. Decolourization of Methyl Orange from fabric by M53 isolate (at 1000mg/L dye concentration, pH 7.0, 37°C under static condition after 21 hours of incubation)

## Discussion

The objective of the current research was to isolate possible azo dye degraders/decolourizers from the rhizospheric and non-rhizospheric soil of the mangrove forest around the Kamothe creek in Navi Mumbai. Azo dyes, which are commonly used in the textile industry, are known to be hazardous to plants and animals (Sudha et al. 2014).

### Effect of dye concentration on decolourization

The dye concentration is thought to be a key factor that influences an organism’s decolorization efficiency. The dye has an inhibiting impact on azo bond breakdown at high concentrations, resulting in less decolorization **(Fig S3)**. This is due to the dye molecule’s toxicity to bacterial cells via limiting metabolic activity, saturation of cells with dye products, electron failure to reach the azo bond chromophore, transport system inactivation, or obstruction of active sites in the Azoreductase enzyme by the dye molecule (Sponza and Isik 2002; Pearce et al. 2006; Vijaykumar et al. 2007). To evaluate the tolerance level and dye decolorization capacity of all isolated bacterial strains, two dye concentration viz. 1000mg and 2000mg/L of MO dye were used. All bacterial strains have achieved the highest decolorization with a dye concentration of 1000 mg/L. **(Fig S3B)**. When the dye concentration was increased from 1000mg/L to 2000mg/L, however, the dye decolourization effectiveness decreased. The dye decolorization was inversely related to dye concentration, which is owing to the inhibitory impact of high dye concentration in the culture media (Singh et al. 2005; Kalme et al. 2007; Vijaykumar et al. 2007; Mohandass and Bhaskar 2008; Ozdemir et al. 2008; Telke et al. 2009). A decline in dye decolorization efficiency with increasing dye concentration may be attributed to the rising harmful effect of dye and its degradation metabolites, which reduces the organisms’ overall efficiency. As a result, the remainder of the investigation was carried out at a constant dye dosage of 1000mg/L.

### Effect of media on dye decolourization

Previous research has demonstrated that increasing the carbon source and changing both the carbon and nitrogen sources causes an increase or decrease in the rate of decolorization. Possibly NB medium contains a complex source of carbon and nitrogen which leads to reduced decolourization of MO by many of the isolates (Aruna et al. 2015). The addition of glucose revealed that all strains were able to use dyes with a high %age of decolorization, which aided bacterial development and, as a result, increased the %age of decolorization rate (Mohan et al. 2002). According to studies by Aruna et al. (2015) the addition of peptone reduced dye decolorization by 20%. As a result, more research into the effects of beef extract, yeast extract, peptone, and other organic nitrogen sources will offer light on why the NB medium shows less or no decolorization. The effect of different carbon and nitrogen sources, as well as the amount of these nutrients, on dye decolorization property was not addressed in the study.

### Effect of cell and Cell Free Extract on dye decolourization

On dye decolorization, the effect of bacterial culture vs. cell free extract was also studied. After 24 hours, CFE is added and the extent of decolorization was monitored for 48-72 hours to compare the extent of decolorization with the culture. This is owing to the presence of enzymes such as Azoreductase, Laccase, and Peroxidase in the cell free extract, which aid in rapid dye decolorization. It’s necessary to investigate the role and significance of enzymes involved in dye decolorization.

### Effect of incubation time on dye decolourization

Because the growth and metabolic rate of different bacteria changes with time, biological decolorization of dye is strongly reliant on the incubation duration. The gradual decolorization of MO dye was observed within the first 30 hours of adding CFE, and then the maximum decolorization was observed within 31-66 hours of incubation, which may contribute to the log phase when rapid metabolic rate and growth of microorganisms aided in the rapid reduction of dye chromophore. The variation in %age decolorization for MO dye could be attributed to physiological differences across all isolated samples. Maximum decolorization was achieved in all samples after 66 hours of incubation.

### Effect of pH on dye decolourization

The pH tolerance of decolorizing isolates was tested in order to see if they might be used in the textile industry. Under alkaline conditions, most reactive azo dyes bind to fibre (cotton) via addition or substitution mechanisms. Below the optimum pH, H^+^ ions effectively compete with dye cations, resulting in a reduction in colour removal efficiency. The highest decolorization of MO dye occurred after 66 hours of incubation at 37°C at pH 7.0 At the optimum pH, the surface of biomass becomes negatively charged, which improves the binding of positively charged dye. Electrostatic attraction causes binding, resulting in a considerable increase in colour removal (Daneshvar et al. 2007). The decolorization of MO dye was reduced as the pH increased. Because the azo bonds will have deprotonated to negatively charged compounds at alkaline pH, azo dye decolorization will be hampered. In acidic pH, the azo bond will protonate, resulting in less decolorization due to a change in chemical structure (Hsueh and Chen 2007). MO dye decolorization increased as pH increased from 6 (4%) to 7 (100%). Hydrogen ions had a significant impact on the metabolic activities of bacteria. At pH 7.0, all isolated bacterial strains showed good decolorization. Decolorization efficiency was shown to be significantly reduced in higher acidic and alkaline conditions because extreme pH environments hinder bacterial development.

### Effect of temperature on dye decolourization

Incubation temperature plays an important role in microbial growth and activity. It is one of the vital parameters taken into consideration for optimization of any kind of bioremediation process. Reduced colour removal below 37°C may be due to the loss of cell viability or thermal non-activation of decolourizing enzymes of isolated bacteria (Panswad and Luangdilok 2000; Çetin D Dönmez G 2006). Decreased decolourization was exhibited at room temperature under static conditions possibly due to the bacterium’s poor growth rate. It implies that the bacterium is mesophilic and possible reason is that the enzyme responsible for decolourization has its activity between 30-40°C (Saratale et al. 2011). Previous report indicates the rapid decolourization of Remazol Black B, Direct Red 81, Acid orange10, Disperse Blue 79, Navy Blue HER and Acid Blue 113 were observed at 37°C (Meehan et al. 2000; Junnarkar et al. 2006; Kolekar et al. 2008). Bacterial species like *Pseudomonas spp*. ETL-1982 also decolourizes the MO dye at 37°C (Shah et al. 2013). Bacterial species like *Enterococcus Sp*, showed that maximum decolourization of sulphonated diazo dye C.I. Direct Red 81 in the temperature range of 30-40°C (Sahasrabudhe et al. 2014). Reports are also available on the complete degradation and decolourization of sulfonated dye (MO) by *Bacillus stratosphericus* SCA1007 without any toxic effect of degraded product (Akansha et al. 2019). A report available on the decolourization of MO dye in the temperature range of 30-40°C (Ahmad Ali Pourbabaee, Malekzadeh Fereydon 2005) which supports our finding.

### Decolourization of mixture of azo dyes using single isolate

The selected bacterial isolate M53 was further used to study the decolorization of a mixture of three dyes (MO, Methyl Red, and Congo Red). This case also demonstrated that a single bacterium was a potent dye decolourizer, with the greatest dye decolorization (100%) attained after 48 hours of incubation at 37°C under static conditions.

According to the above interpretation, the selected bacterium is tolerable to all three azo dyes, and this bacterial isolate can be exploited for the degradation of azo dyes in industries. More research is needed to determine the precise methods and nature of dye degradation.

### FTIR analysis of decolourized product of dye

FTIR analysis of treated and untreated dye with medium at 48/72 hours was compared. The FTIR pattern of the decolorized product lacks a peak at 1597.1 cm^-1^, indicating the disappearance of the (-N=N-) azo bond (Telke et al. 2009). Telke et al. (2009) found comparable results for the removal of peaks during the research of biochemical characterisation and potential for textile dye degradation of *Aspergillus ochraceus* NCIM-1146 Blue Laccase. This suggests that the dye was degraded and showed new peaks, thus more research is needed to determine the effect of incubation time on seeing these new peaks (PS et al. 2013).

### Phylogenetic analysis of 16S rRNA sequence

The 16S rRNA sequence analysis is employed as a framework for the modern classification of bacteria. In the present study 7 isolates A1, A2, M41, M42, M43, M3 and M53 were identified as *Bacillus paramycoides, Pseudomonas taiwanensis, Acinetobacter pitti, Exiguobacterium acetylicum, Psychrobacter celer, Citrobacter murliniae* and *Aeromonas taiwanensis* for decolourization of MO dye. There are some previous reports that some strains of *Pseudomonas* sp. are able to decolourize the number of toxic compounds. Reports are available on dye decolourization of MO dye by *Pseudomonas* sp. isolated from textile effluent (Maulin P Shah et al 2013). Recent findings by (Chaudhary et al. 2021) indicated the beneficial effect of the organism *Pseudomonas taiwanensis* as PGPR. In her studies the combined application of nanogypsum and *Pseudomonas taiwanensis* augmented the growth of maize by improving the structure and function of soil without causing any toxic effect. The bacterial strain *Aeromonas hydrophila* and *P. monteilii* isolated from the dye industry area were selected for the decolourization of MO dye and Methylene Blue dye respectively (Fulekar et al. 2013). The bacterial strain *Acinetobacter radioresistens* isolated from the contaminated area of textile industry was selected for the decolourization of Acid Red dye (Ramya M et al. 2010)

### Application based study for dye decolourization

Azo dyes are coloured chemicals that are used to colour textiles, silk, wool, food, and other items. When azo dye is applied to fabrics, it forms a hydrogen bond with the dying process. In this investigation, MO dye was used to dye cloth, and the coloured fabric was then introduced in a MBM media containing CFE of M53 isolate. The results of this study reveal that after 21 hours of incubation at 37°C in a static condition, the fabric had completely decolourized.

## Conclusion

We propose that these isolated bacterial cultures will play an important role in mitigating the anthropogenic environmental disaster because this saline wetland of Navi Mumbai, which includes mangrove forest, is undergoing a lot of climatic change in saline content, drought and wet condition temperature variation, and oxygen. As a result, these microorganisms and their enzymes will have greater potential in various bioremediation processes, as well as bio fertiliser applications for supporting plant development, soil health, and the environment. This research suggests that all of the possible dye decolourizers discovered in this area might be used to decolorize and degrade azo dyes as a more environmentally friendly and cost-effective alternative to physicochemical decomposition procedures. Further research into the effects of environmental variables such as pH, temperature, incubation time, dyes and their concentrations, and the effects of various abiotic stresses on dye decolorization rates and their role as PGPR would benefit the agricultural and environmental communities.

## Abbreviations

MBM: Minimal Basal Medium
CFE: Cell Free Extract
MO: Methyl Orange
MR: Methyl Red
CR: Congo Red

## Acknowledgement

The author is grateful to the D Y Patil Deemed To Be University’s School of Biotechnology and Bioinformatics in Navi Mumbai, India, for providing research facilities for this study.

## Declaration

The authors declare that no funds, grants, or other support were received during the preparation of this manuscript.”

## Author Contribution

AM: Experiment perform, Compilation of data, Manuscript drafting, Graphical abstract preparation SS: Inception of the project, planning and execution of experiment, Manuscript editing and reviewing. NP: Phylogenetic analysis, collection of soil sample

JP: Inception of the project, planning of the project, Collection of soil samples

## References

Abbas SZ, Yee CJ, Hossain K, et al (2019) Isolation and characterization of mercury-resistant bacteria from industrial wastewater. Desalin Water Treat 138:128–133. https://doi.org/10.5004/dwt.2019.23279

Ahmad Ali Pourbabaee, Malekzadeh Fereydon SMN and MA (2005) Decolorization of Methyl Orange (As a Model Azo Dye) by the Newly Discovered Bacillus Sp. Iran J Chem \& Chem Eng English Ed 24:41–45. https://doi.org/10.1002/9783527611904.ch87

Akansha K, Chakraborty D, Sachan SG (2019) Decolorization and degradation of methyl orange by Bacillus stratosphericus SCA1007. Biocatal Agric Biotechnol 18:101044. https://doi.org/10.1016/j.bcab.2019.101044

Aruna B, Silviya LR, Kumar ES, et al (2015) Original Research Article Decolorization of Acid Blue 25 dye by individual and mixed bacterial consortium isolated from textile effluents. Int J Curr Microbiol Appl Sci 4:1015–1024

Carliell CM, Barclay SJ, Naidoo N, et al (1995) Microbial decolourisation of a reactive azo dye under anaerobic conditions. ISSN 0378-4738=Water SA 21:61–69

Çetin D Dönmez G (2006) Decolorization of reactive dyes by mixed cultures isolated from textile effluent under anaerobic conditions Demet C. Enzyme Microb Technol 38:926–930. https://doi.org/10.1016/j.enzmictec.2005.08.020

Chaudhary P, Khati P, Chaudhary A, Maithani D (2021) Cultivable and metagenomic approach to study the combined impact of nanogypsum and Pseudomonas taiwanensis on maize plant health and its rhizospheric microbiome. PLoS One 1–17. http://dx.doi.org/10.1371/journal.pone.0250574

Chen KC, Wu JY, Liou DJ, Hwang SCJ (2003) Decolorization of the textile dyes by newly isolated bacterial strains. J Biotechnol 101:57–68. https://doi.org/10.1016/S0168-1656(02)00303-6

Chiong T, Yon S, Hong Z, et al (2016) Journal of Environmental Chemical Engineering Enzymatic treatment of methyl orange dye in synthetic wastewater by plant-based peroxidase enzymes. Biochem Pharmacol 4:2500–2509. https://doi.org/10.1016/j.jece.2016.04.030

D.T. Sponza Mi (2002) Decolorization and azo dye degradation by anaerobic / aerobic sequential process. Enzyme Microb Technol 31:102–110. https://doi.org/10.1016/S0141-0229(02)00081-9

Daneshvar N, Ayazloo M, Khataee AR, Pourhassan M (2007) Biological decolorization of dye solution containing Malachite Green by microalgae Cosmarium sp. 98:1176–1182. https://doi.org/10.1016/j.biortech.2006.05.025

Felsenstein J (1985) Confidence limits on phylogenies: an approach using the bootstrap. Int J Org Evol 39:783–791. https://doi.org/10.1111/j.1558-5646.1985.tb00420.x

Forgacs E, Cserháti T, Oros G (2004) Removal of synthetic dyes from wastewaters: A review. Environ Int 30:953–971. https://doi.org/10.1016/j.envint.2004.02.001

Fulekar MH, Wadgaonkar SL, Singh A (2013) Decolourization of Dye Compounds by Selected Bacterial Strains isolated from Dyestuff Industrial Area. Int J Adv Res Technol 2:182–192

Guo G, Liu C, Hao J, et al (2021) Development and characterization of a halo-thermophilic bacterial consortium for decolorization of azo dye. Chemosphere 272:129916. https://doi.org/10.1016/j.chemosphere.2021.129916

Heer and Somesh (2017) MICROBIAL PIGMENTS AS A NATURAL COLOR: A REVIEW Kanchan Heer *1 and Somesh Sharma 2 Department of Biotechnology *1, Department of Bioengineering and Food Technology 2, Shoolini University of Biotechnology and Management Sciences, Bhajhol, Solan, Himacha. IJPSR 8:1913–1922. https://doi.org/10.13040/IJPSR.0975-8232.8(5).1913-22

Hsueh C, Chen B (2007) Comparative study on reaction selectivity of azo dye decolorization by Pseudomonas luteola. J Hazard Mater 141:842–849. https://doi.org/10.1016/j.jhazmat.2006.07.056

Junnarkar N, Murty DS, Bhatt NS, Madamwar D (2006) Decolorization of diazo dye Direct Red 81 by a novel bacterial consortium. World J Microbiol Biotechnol 163–168. https://doi.org/10.1007/s11274-005-9014-3

Kalme SD, Parshetti GK, Jadhav SU, Govindwar SP (2007) Biodegradation of benzidine based dye Direct Blue-6 by Pseudomonas desmolyticum NCIM 2112. Bioresour Technol 98:1405–1410. https://doi.org/10.1016/j.biortech.2006.05.023

Kolekar YM, Pawar SP, Gawai KR, et al (2008) Bioresource Technology Decolorization and degradation of Disperse Blue 79 and Acid Orange 10, by Bacillus fusiformis KMK5 isolated from the textile dye contaminated soil. Bioresour Technol 99:8999–9003. https://doi.org/10.1016/j.biortech.2008.04.073

Lalnunhlimi S, Veenagayathri K (2016) Decolorization of azo dyes (Direct Blue 151 and Direct Red 31) by moderately alkaliphilic bacterial consortium. Brazilian J Microbiol 47:39–46. https://doi.org/10.1016/j.bjm.2015.11.013

Letunic I, Bork P, Gmbh BS (2021) Interactive Tree Of Life (iTOL) v5: an online tool for phylogenetic tree display and annotation. Nucleic Acids Res 49:293–296. https://doi.org/10.1093/nar/gkab301

Maulin P Shah, Kavita A Patel AMD (2013) Microbial Degradation and Decolorization of Methyl Orange Dye by an Application of Pseudomonas Spp. Int J Environ Bioremediation Biodegrad 1:26–36. https://doi.org/10.12691/ijebb-1-1-5

Meehan C, Banat IM, Mcmullan G, et al (2000) Decolorization of Remazol Black-B using a thermotolerant yeast, Kluyveromyces marxianus IMB3. Environ Int 26:75–79. https://doi.org/10.1016/S0160-4120(00)00084-2

Mohan SV, Rao NC, Prasad KK, Karthikeyan J (2002) Treatment of simulated Reactive Yellow 22 (Azo) dye effluents using Spirogyra species. Waste Manag 22:575–582. https://doi.org/10.1016/S0956-053X(02)00030-2

Mohandass R, Bhaskar A (2008) Decolorization and biodegradation of Indigo carmine by a textile soil isolate Paenibacillus larva Decolorization and biodegradation of Indigo carmine by a textile soil isolate Paenibacillus larvae. Biodegradation 19:283–291. https://doi.org/10.1007/s10532-007-9134-6

Morgulis A, Coulouris G, Raytselis Y, et al (2008) Database indexing for production MegaBLAST searches. 24:1757–1764. https://doi.org/10.1093/bioinformatics/btn322

Ozdemir G, Pazarbasi ÆB, Kocyigit ÆA, Ersoy ÆE (2008) Decolorization of Acid Black 210 by Vibrio harveyi TEMS1 a Newly Isolated Bioluminescent Bacterium from Izmir Bay, Turkey. World J Microbiol Biotechnol 24:1375–1381. https://doi.org/10.1007/s11274-007-9619-9

Panswad T, Luangdilok W (2000) DECOLORIZATION OF REACTIVE DYES WITH DIFFERENT MOLECULAR STRUCTURES UNDER DIFFERENT ENVIRONMENTAL CONDITIONS. Water Res 34:4177–4184. https://doi.org/10.1016/S0043-1354(00)00200-1

Pearce CI, Christie R, Boothman C, et al (2006) Reactive Azo Dye Reduction by Shewanella Strain J18 143. Biotechnol Bioeng 95:692–73. https://doi.org/10.1002/bit.21021

Pearce CI, Lloyd JR, Guthrie JT (2003) The removal of colour from textile wastewater using whole bacterial cells: A review. Dye Pigment 58:179–196. https://doi.org/10.1016/S0143-7208(03)00064-0

Ps H, Joseph L A D, (2013) Photocatalytic degradation of textile dyes by hydrogel supported titanium dioxide nanoparticles. J Environ Eng Ecol Sci 2:2. https://doi.org/10.7243/2050-1323-2-2

Ramya M, Iyappan S MA and JJS (2010) Biodegradation and Decolorization of Acid Red by Acinetobacter radioresistens. J Bioremediation Biodegrad 01:1–6. https://doi.org/10.4172/2155-6199.1000105

Sahasrabudhe MM, Saratale RG, Saratale GD, Pathade GR (2014) Decolorization and detoxification of sulfonated toxic diazo dye C. I. Direct Red 81 by Enterococcus faecalis YZ 66. J Environ Heal Sci Eng 12:1–13. https://doi.org/10.1186/s40201-014-0151-1

Saitou N, Nei M (1987) The Neighbor-joining Method: A New Method for Reconstructing Phylogenetic Trees ‘. Mol Biol Evol 4:406–425. https://doi.org/10.1093/oxfordjournals.molbev.a040454

Saratale RG, Saratale GD, Chang JS, Govindwar SP (2011) Bacterial decolorization and degradation of azo dyes: A review. J. Taiwan Inst. Chem. Eng. 42:138–157

Saravanan R, Manoj D, Qin J, et al (2018) Mechanothermal synthesis of Ag/TiO2 for photocatalytic methyl orange degradation and hydrogen production. Process Saf Environ Prot 120:339–347. https://doi.org/10.1016/j.psep.2018.09.015

Shah MP, Ka P, Ss N, Am D (2013) Bioremoval of Azo dye Reactive Red by Bacillus spp. ETL-1982 Bior emediation & Biodegradation. 4:3–7. https://doi.org/10.4172/2155-6199.1000186

Shaw CB, Carliell CM, Wheatley AD (2002) Anaerobic/aerobic treatment of coloured textile effluents using sequencing batch reactors. Water Res 36:1993–2001. https://doi.org/10.1016/S0043-1354(01)00392-X

Singh K, Arora S (2011) Removal of synthetic textile dyes from wastewaters: A critical review on present treatment technologies. Crit Rev Environ Sci Technol 41:807–878. https://doi.org/10.1080/10643380903218376

Singh M, Singh H, Kumar D, et al (2005) Decolorization of various azo dyes by bacterial consortium. Dye Pigment 67:55–61. https://doi.org/10.1016/j.dyepig.2004.10.008

Singh RP, Singh PK, Singh RL (2017) Role of Azoreductases in Bacterial Decolorization of Azo Dyes. 9:9–11. https://doi.org/10.19080/CTBEB.2017.09.555764

Sudha M, Saranya A, Selvakumar G, Sivakumar N (2014) Microbial degradation of Azo Dyes: A review. IntJCurrMicrobiolAppSci 3:670–690

Tamura K, Stecher G, Kumar S (2021) MEGA11: Molecular Evolutionary Genetics Analysis Version 11. 38:3022–3027. https://doi.org/10.1093/molbev/msab120

Telke AA, Kalyani DC, Dawkar V V, Govindwar SP (2009) Influence of organic and inorganic compounds on oxidoreductive decolorization of sulfonated azo dye C. I. Reactive Orange 16. 172:298–309. https://doi.org/10.1016/j.jhazmat.2009.07.008

Vijaykumar MH, Vaishampayan PA, Shouche YS, Karegoudar TB (2007) Decolourization of naphthalene-containing sulfonated azo dyes by Kerstersia sp. strain VKY1. 40:204–211. https://doi.org/10.1016/j.enzmictec.2006.04.001

Weisburger JH (2002) Comments on the history and importance of aromatic and heterocyclic amines in public health. Mutat Res - Fundam Mol Mech Mutagen 506–507:9–20. https://doi.org/10.1016/S0027-5107(02)00147-1

Yoon S, Ha S, Kwon S, et al (2017) Introducing EzBioCloud: a taxonomically united database of 16S rRNA gene sequences and whole-genome assemblies. Int J Syst Evol Microbiol 1613–1617. https://doi.org/10.1099/ijsem.0.001755

Zollinger H, Morgan TH, The M (2003) Color chemistry: syntheses, properties, and applications of organic dyes and pigments. John Wiley & Sons

