## Supplementary data for "Screening and Identification of Azo Dye Decolourizers from Mangrove Rhizospheric soil"

**Supplementary file**

**S1. Analysis of bacterial growth**

The growth of all isolated bacterial cultures was spectrophotometrically monitored at 600nm at varied time intervals during dye decolorization.

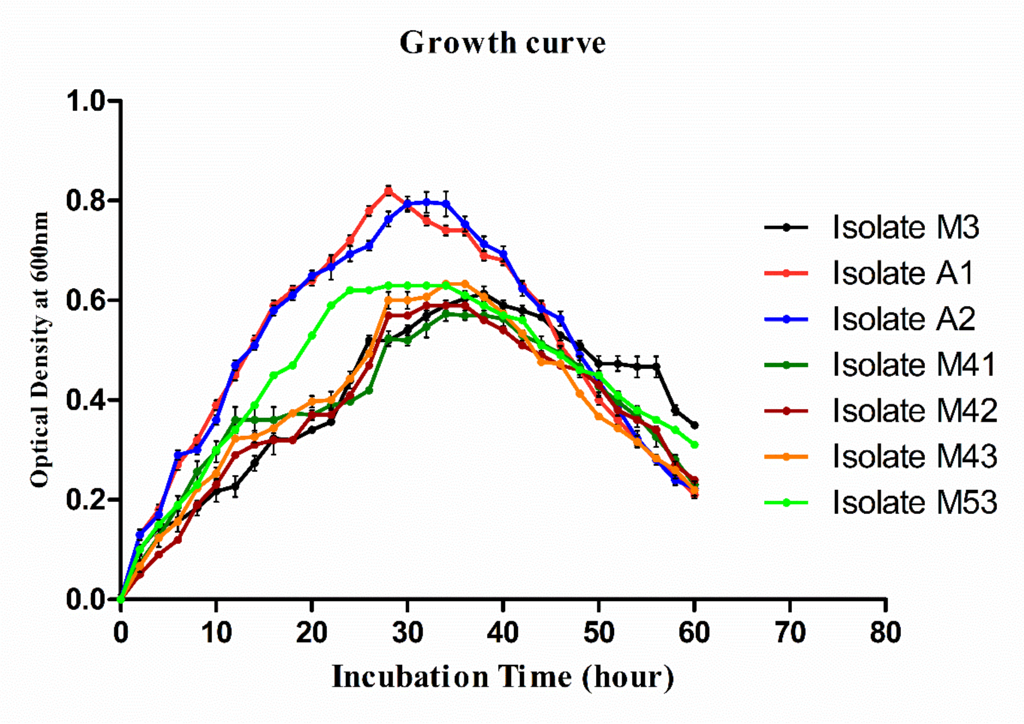


**Figure S1: Growth curve of all bacterial isolates during dye decolourization**

**S2. Screening of dye decolourizers based on clear zone on Nutrient agar plate containing Methyl Orange dye.**

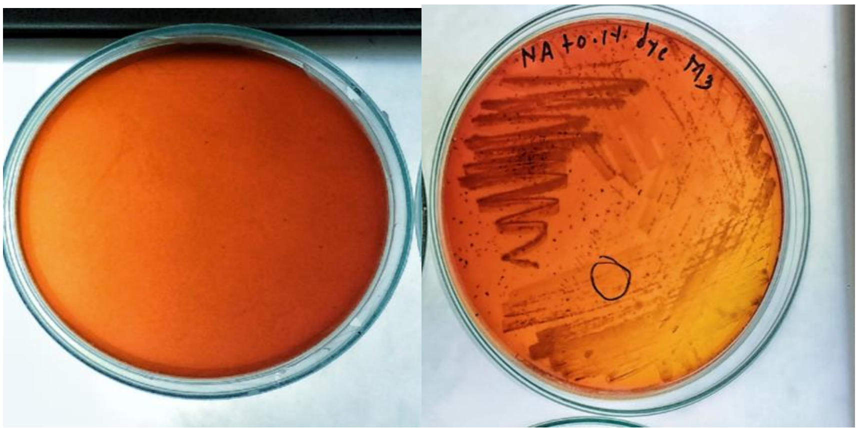


**a- control NA plate with 0.1% MO dye   b- decolourization zone on NA plate with 0.1% MO**

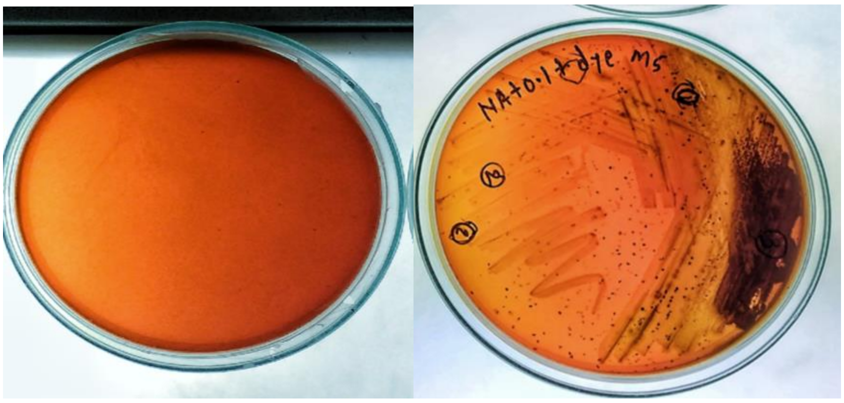


**c- control NA plate with 0.1% MO dye   d- decolourization zone on NA plate with 0.1% Mo**

**Figure S2. Bacterial Colonies from five soil sample showing decolourization zone on NA plate with 0.1% dye.**

**S3. Effect of dye concentration on dye decolourization**

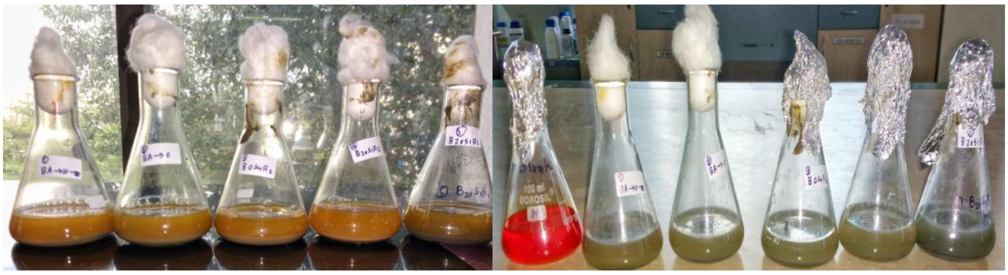

**a- At zero-hour incubation in NB** **b- At 48-hour incubation in NB**

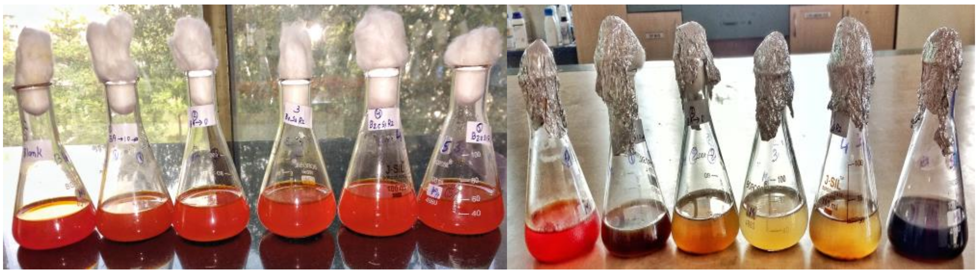

**c- At zero-hour incubation in MBM** **d-At 48-hour incubation in MBM**

**Figure S3A. Enrichment of five soil sample in NB media (a&b) and (c&d) MBM media with 0.2% Methyl Orange at pH 7.0, 25°C under static condition**

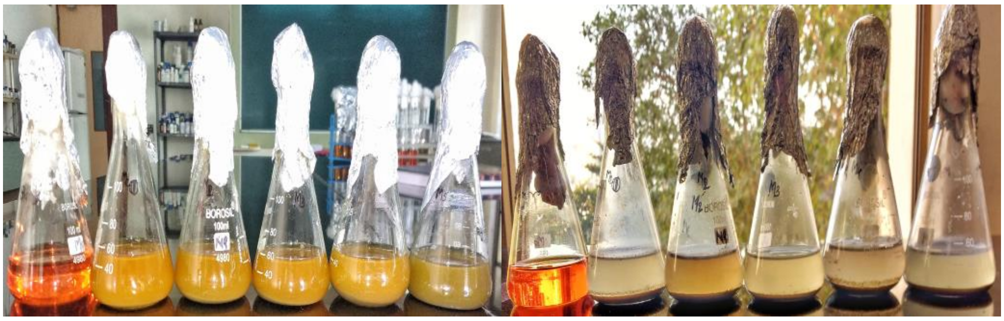

**a-At zero-hour incubation in MBM** **b-At 48-hour incubation in MBM**

**Figure S3B. Enrichment of five soil sample in MBM media with 0.1% Methyl Orange at pH 7.0, 25°C under static condition**

**S4. Effect of media on dye decolourization**

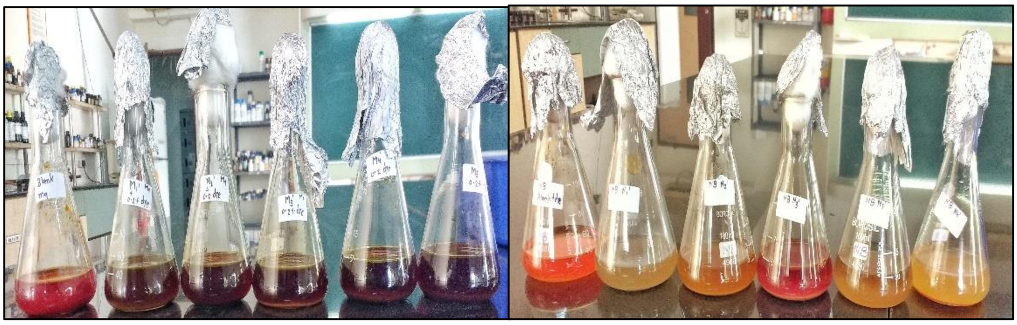

**Figure S4. Effect of media on dye decolourization of Methyl Orange**

**S5. Effect of pH on dyed decolourization.**

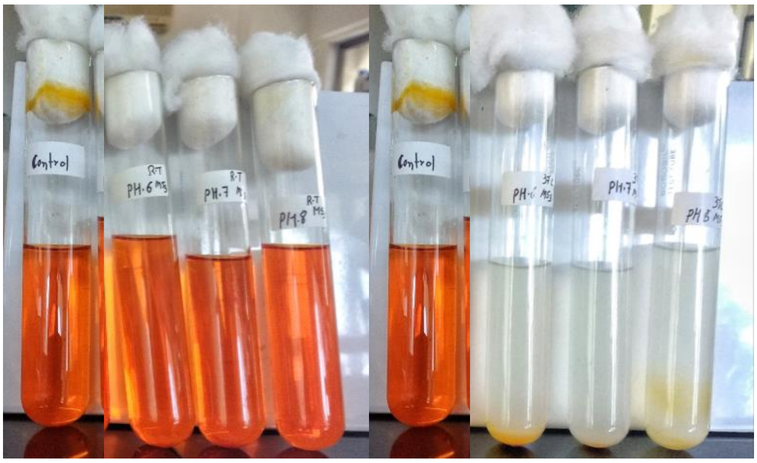

**a-At zero-hour incubation b-At 26-hour incubation**

**Figure S5. Effect of pH on dye decolourization of Methyl Orange (at 1000 mg/L dye concentration, 37°C under static.**

**S6. Effect of temperature on dye decolourization**

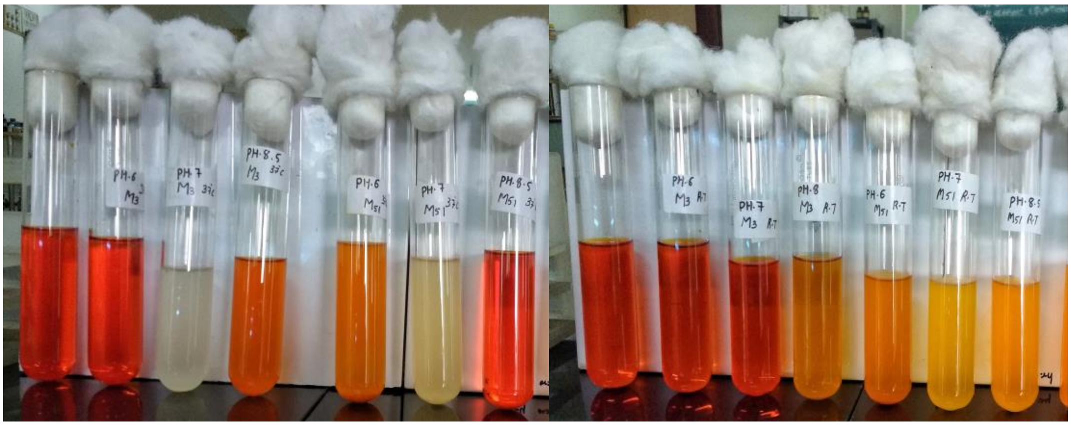

**a- Decolourization at 37°C** **b- Decolourization at RT**

**Figure S6. Effect of temperature on decolourization of Methyl Orange (at 1000 mg/L dye concentration, pH 7.0, under static condition after 66 hours)**

**Table S1. Summary of the closest neighbours of 7 strains from Kamothe area**

| Isolate No. | Closest Neighbour | Accession No. | % Similarity |
|---|---|---|---|
| **A1** | *Bacillus Paramycoides* NH24A2(T) | MT256266 | 100% |
| **A2** | *Pseudomonas taiwanensis* BCRC 1775 | MT256267 | 100% |
| **M41** | *Citrobacter murliniae CDC 2970-59(T)* | MT256269 | 99.12% |
| **M42** | *Acinetobacter pitti* CIP 70.29 | MT256270 | 99.90% |
| **M43** | *Exiguobacterium acetylicum* DSM 20416 | MT256271 | 99.53% |
| **M3** | *Psychrobacter celer* SW-238 | MT256273 | 99.17% |
| **M53** | *Aeromonas taiwanensis* A2-50 | MT256274 | 99.88% |

**Table S2. One-way ANOVA analysis of 7 isolates at different pH (6, 7, and 8.5) and different temp (25°C and 37°C) after 66 hours**

| One-way ANOVA analysis of 7 isolates at different pH and different temperature | | | | | | | |
|---|---|---|---|---|---|---|---|
| | **Source of Variation** | **SS** | **df** | **MS** | **F** | **P-value** | **F crit** |
| **7 isolates at pH range of (6, 7, and 8.5) at 37°C** | pH 6.0 | 3.033431 | 7 | 0.202229 | 2311.186 | 3.73E-44 | 1.99199 |
| | pH 7.0 | 3.253748 | 7 | 0.216917 | 2539.511 | 8.27E-45 | 1.99199 |
| | pH 8.5 | 4.175315 | 7 | 0.278354 | 7859.416 | 1.18E-52 | 1.99199 |
| **7 isolates at 25°C and 37°C at pH 7.0** | Temp 37°C | 3.253748 | 7 | 0.216917 | 2539.511 | 8.27E-45 | 1.99199 |
| | Temp 25°C | 4.26313 | 7 | 0.284209 | 2236.393 | 6.3E-44 | 1.99199 |
